# New insights into tissue-specific responses and interactive characteristics of crop-microbe “One Health” system to soil chromium and ofloxacin pollution

**DOI:** 10.1101/2023.10.08.561380

**Authors:** Jia-Min Xu, Yaru Zhang, Kai Wang, Guodong Zhang, Ying Liu, Hao-Ran Xu, Hu-Yi Zi, Ai-Jie Wang, Hao-Yi Cheng, Yao Lv, Kun Xu

## Abstract

This study firstly investigated the tissue-specific responses and interactive characteristics of the crop-microbe system to co-pollution with ofloxacin (OFL) and chromium (Cr) in soil. The results emphasized the hormesis effect induced by low-dose OFL (1 mg L^-1^) on ginger plants subjected to soil Cr stress. However, high-dose OFL (100 mg L^-1^) and Cr co-stressed plants exhibited reduced growth, root activity, antioxidant enzyme activities and photosynthesis-fluorescence performances, while the reactive oxygen species (ROS) reflected by O_2_^·-^ and H_2_O_2_ significantly increased up to 43.34% and 78.63%, respectively, compared to other treatments. In addition, high-throughput sequencing indicated that OFL influenced rhizosphere microbial diversity, composition, and evolution, favoring *Proteobacteria* proliferation under co-pollution. Meanwhile, root exudate patterns shifted, with humic-like exudates, potentially interacting with pollutants and microbes. Notably, enrichments of antibiotic resistance gene (*qnr*S) in edible rhizome and potential pathogenic bacteria in ginger rhizosphere were observed under OFL and Cr co-pollution, raising environmental and food chain concerns. Through structural equation modeling, we quantitatively established correlations within soil-crop-microbe factors, emphasizing their interconnected nature. Overall, this study sheds light on the complex responses of the crop-microbe system to OFL and Cr co-pollution, highlighting the importance of understanding pollutant interactions for enhancing plant resilience and mitigating environmental risks.

**Highlights:**

- Revealed a hormesis from low-dose OFL on Cr-polluted ginger, enhancing resilience.
- Rhizosphere microbial composition shift under co-pollution, favoring Proteobacteria.
- *qnr*S and potential pathogens increased under co-pollution, posing food chain risks.
- Structural equation modeling quantified correlations in soil- crop-microbe factors.
- Clarified crop-microbe “One Health” interactions’ role under Cr & OFL co-pollution.

**Graphical abstract:** 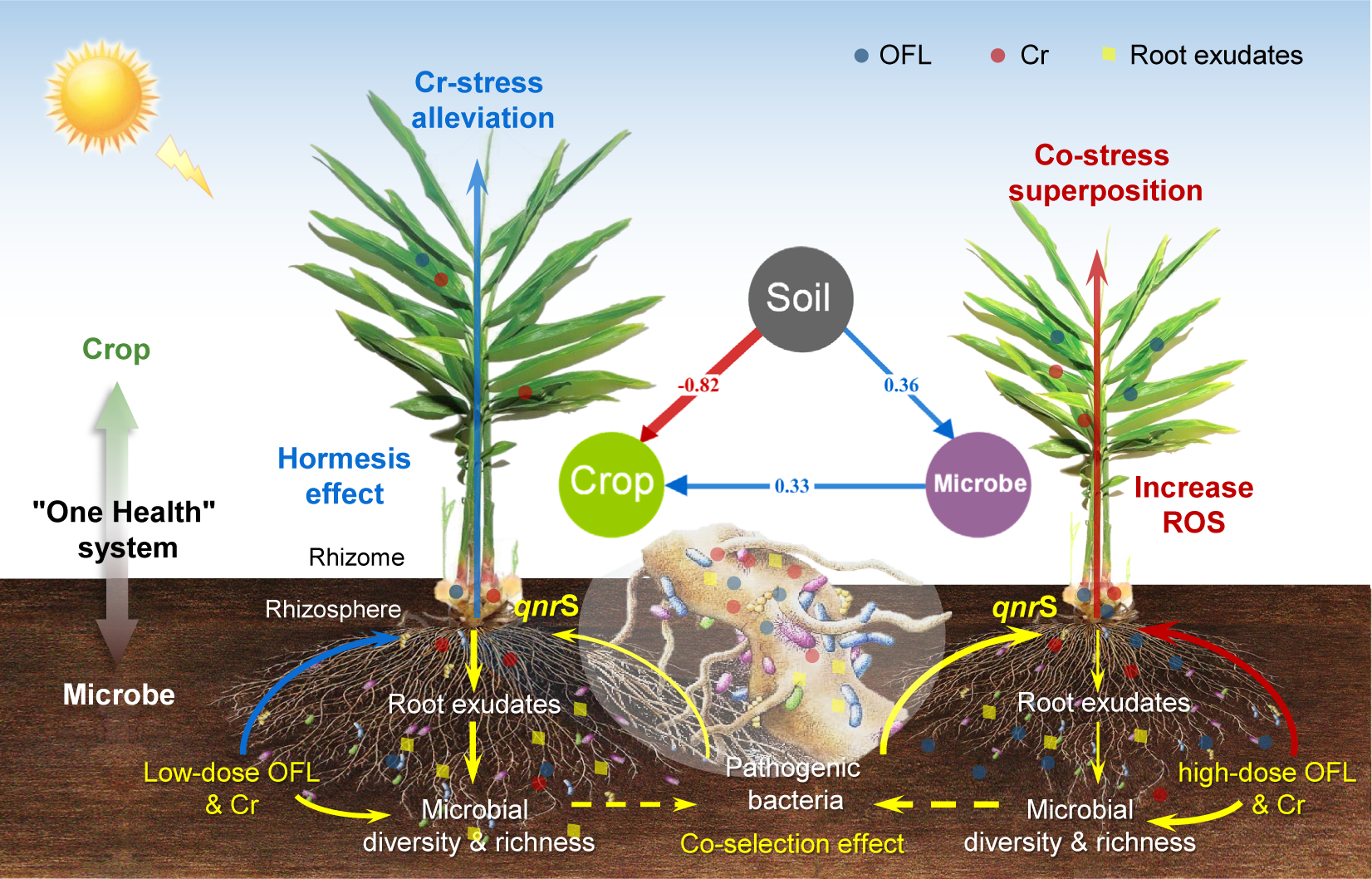

## 1 Introduction

The escalating global utilization of antibiotics has reached an unprecedented scale, with an estimated annual consumption exceeding 42 million kg (Klein et al., 2018). This widespread usage of antibiotics has raised profound concerns regarding the emergence of antibiotic resistance genes (ARGs), which poses a critical threat to various aspects of agriculture, including crop production, food safety, and human health (WHO 2017; Yin et al., 2023). In agroecosystem, heavy metals are ubiquitous pollutants originating from industrial activities and natural sources, which presence can influence the fate of antibiotics by altering soil physicochemical properties, microbial communities, and metabolic processes (Lyu et al., 2022). To be noted, the co-occurrence of heavy metals and antibiotics can synergistically affect the mobility of both pollutants, potentially exacerbating their environmental and human health risks (Xu et al., 2023). Furthermore, the selection pressure imposed by heavy metal contamination might enhance the horizontal transfer of ARGs among microorganisms, further contributing to the global challenge of antibiotic resistance development (Di Cesare et al., 2016). Thus, environmental pollution caused by heavy metals and antibiotics has become a critical concern globally, impacting agroecological system and human well-being.

Chromium (Cr) and ofloxacin (OFL) are typical pervasive pollutants of Heavy metals and antibiotics in agroecosystem respectively (Lv et al., 2020), which pose distinct yet interrelated challenges. Cr, often introduced into soil through industrial activities and agricultural runoff, poses threats to soil quality, groundwater, and ecosystem (Kushwaha and Singh, 2020; Prasad et al., 2021). High Cr levels can impede root elongation, disrupt nutrient uptake, and lead to oxidative stress in plants (Guo, 2021; Tripathi et al., 2017). Simultaneously, OFL, as a fluoroquinolone antibiotic and widely used in veterinary and human medicine, can accumulate in soil and thus profoundly disrupt the delicate equilibrium of the crop-microbe health continuum (Luo et al., 2010; Zhu et al., 2013). Currently, the presence of OFL in soil environments has been linked to altered microbial diversity, shifts in functional profiles, and the proliferation of ARGs among microbial communities (Lv et al., 2020). These alterations not only compromise soil health but also have far-reaching implications for ecosystem stability and sustainability. Understanding the intricate interactions between heavy metals and antibiotics is essential for devising effective strategies to mitigate their collective impacts on soil quality, plant health, and ecosystem sustainability.

The concept of “One Health” underscores the interconnectedness of human, animal, and environmental health, emphasizing the need for integrated approaches to tackle multifaceted challenges. In the context of soil pollution, the crop-microbe “One Health” system exemplifies this interdependence, where soil microorganisms and crops together constitute a complex web of interactions that shape agricultural productivity (e.g., soil fertility, plant health, and food security) and environmental health (Lyu et al., 2022). As known, microorganisms play a vital role in soil ecosystems, contributing to nutrient cycling, organic matter decomposition, and pollutant remediation (Wang et al., 2021). Their diverse metabolic capacities can transform and mitigate pollutants, but the combined impact of multiple pollutants on microbial communities remains poorly understood. Studies have shown that microbial communities can exhibit varying sensitivities to different pollutants, leading to shifts in community composition and functional diversity (Ding et al., 2021; Zhao et al., 2017). Since interactions between microorganisms and pollutants can lead to unforeseen effects, necessitating a comprehensive assessment of microbial responses to dual contamination. In addition, crops, as integral components of the “One Health” system, are directly affected by soil pollutants due to their intimate connection to the soil environment. The potential accumulation of heavy metals and antibiotics in crops raises concerns about food safety and quality, as well as the impact on human health. While the individual effects of pollutants on crops have been extensively studied (Lyu et al., 2022; J. Xu et al., 2023), investigations into the combined effects of multiple pollutants are limited. Therefore, the concurrent exposure of this crop-microbe “One Health” system to Cr and OFL pollution demands a thorough investigation into tissue-specific responses and interactive characteristics to assess the potential environmental risks and implications for agricultural practices.

Ginger (Zingiber officinale), as a staple agricultural crop with a worldwide distribution and an annual production exceeding 2.5 million tons (FAOSTAT, 2020), holds both economic significance and medicinal value. However, Recent studies have highlighted the potential for ginger rhizomes to accumulate heavy metals and antibiotics from contaminated soil, raising concerns about potential human exposure through consumption (Lv et al., 2020; Xu et al., 2020b). Given its importance and susceptibility to soil pollutants, ginger was selected as an ideal model crop here to investigate the interactive effects of soli OFL and Cr on the crop-microbe “One Health” system. This study aims to fill existing knowledge gaps by: (i) revealing the tissue-specific responses of ginger to simultaneous exposure to soil Cr and OFL pollution; (ii) elucidating the interactive characteristics of the ginger-microbe system under dual pollution scenarios; and (iii) assessing the associated risks to agricultural sustainability and human health. This research will contribute to a more comprehensive understanding of the multifaceted relationships between soil pollutants and crop-microbe system within agroecosystems, which is crucial for devising effective pollution management strategies and promoting sustainable agricultural practices.

## 2 Materials and methods

### 2.1 Experimental approach

The pot experiments of ginger plants were performed at a solar greenhouse in Horticultural Experimental Station of Shandong Agricultural University from May 2020 to November 2020. The ginger cultivar for this experiment was “*Shannong* 1”. Each pot (height: 30 cm, diameter: 25 cm) containing 8 kg of air-dried soil was sown with two ginger plants. The chemical properties of the air-dried soil were examined before the experiment with a pH of 7.45 and alkali-hydrolyzed nitrogen, available phosphorus, and available potassium contents of 100 mg kg^-1^, 64 mg kg^-1^, and 128 mg kg^-1^, respectively. The different pollution solutions were applied evenly into each pot at one time by sewage irrigation (Xu et al., 2020b). Three replicate plot groups were set up for each treatment condition, where a completely randomized block arrangement was used, with five pots per plot. By setting up three different groups of simulated pollution treatments, this work focused on the effect of soil co-pollution with Cr and OFL on the microbial-crop “One Health” system. Specifically, Cr (i.e., single Cr pollution, 100 mg·L^-1^ Cr), (i.e., low dose of OFL and Cr co-pollution, 1 mg·L^-1^ OFL & 100 mg·L^-1^ Cr) and (i.e., high dose of OFL and Cr co-pollution, 100 mg·L^-1^ OFL & 100 mg·L^-1^ Cr), respectively. Here, 100 mg·L^-1^ Cr was chosen as the simulated pollution concentration of heavy metals for two reasons: 1) to be close to the average detection level of Cr in most soils of China, and 2) to be low risk to the safety of most agricultural products and soil ecological environment (Pan et al., 2016; Yang et al., 2018). In addition, 1 mg·L^-1^ and 100 mg·L^-1^ OFL were chosen as the antibiotic pollution concentrations, respectively, since our previous experiments have documented that the former was able to promote the growth of ginger plants while the latter produced an inhibitory effect (Lv et al., 2020).

### 2.2 Analytical methods of crop physiological parameters

Different organs of the ginger plant (i.e., root, leaf, and rhizome) were sampled at the harvest period (about five months after cultivation). The dry weights of the samples were recorded after drying at 75 °C. For the content of antioxidant enzymes in the fresh leaves of the ginger, the superoxide dismutase (SOD), peroxidase (POD) and catalase (CAT) were measured according to the previously reported methods (Prochazkova et al., 2001). Meanwhile, the hydrogen peroxide (H_2_O_2_), superoxide anion (O_2_^·-^) and malondialdehyde (MDA) were determined as the procedures described by Xu et al. (2020a). To evaluate the photosynthetic parameter variations among different pollution groups, the third functional leaves of ginger plant were selected to determine the intercellular CO_2_ concentration (Ci), stomatal conductance (Gs) and net photosynthetic rate (Pn), by employing a CIRAS-3 portable photosynthetic system (PP Systems, USA). Besides, the initial fluorescence (Fo), maximal fluorescence (Fm), variable fluorescence/fluorescence maximum (Fv/Fm), photochemical quantum efficiency of photosystem II (φPSII), nonphotochemical quenching coefficient (NPQ), photochemical quenching coefficient (qP) and the photosynthetically active radiation-photosynthetic electron transfer rate (PAR-ETR) were determined in real time, using a Imaging-PAM chlorophyll fluorometer (HeinzWalz GmbH, Effeltrich, Germany).

For the different pollution groups, one ginger plant with intact root system was taken from each, washed of root attachment and placed in 1 L of pure water for 24 h under dark conditions. To understand the root exudates of the ginger plants, fluorescence excitation emission matrix (EEM) of the water samples filtered through a 0.45 μm cellulose membrane were measured by a fluorescence spectrophotometer (RF-6000, Shimadzu, Japan). Further, the data were further analysed according to parallel factor analysis (PARAFAC). The details can be found in Text S1.

### 2.3 Quantification of ARGs

The ginger rhizome samples were collected at the harvest period. The pretreatment procedures for these samples, DNA extraction and PCR procedures could be found in our previous report (Xu et al., 2020a). The 16S rRNA gene and the typical ofloxacin resistance gene (*qnr*S) were quantified based on the quantitative PCR (qPCR), and the primer pairs used in qPCR assays for quantification could be found in Table S1. Three parallel lines were set up for each sample. q-PCR reaction standard curves had correlation coefficients (*R^2^*) greater than 0.99, with amplification efficiencies all in the range of 90% ∼ 110%, suggesting the validity of the qPCR reactions met the experimental requirements.

### 2.4 Analyzation of rhizosphere microbial community

During the harvest period, the rhizosphere soils of the ginger plant were collected separately and stored at −80 ℃ to be prepared for DNA extraction. Samples were pre-processed according to the previous methods (Beckers et al., 2016). The 16S rRNA V3-V4 variable regions were amplified by polymerase chain reaction (PCR) using the universal primers 338F (5’-ACTCCTACGGGAGGCAGCAG-3’) and 806R (5’-GGACTACHVGGGTWTCTAAT3’). Polymerase chain reaction (PCR) amplification was performed using TransGen AP221-02: TransStart Fastpfu DNA Polymerase; PCRcc instrument: ABI GeneAmp® 9700. The obtained sequences were assembled and assembled into complete fragments, and OTUs (operational taxonomic units) were divided at the 97% sequence similarity level to calculate the relative abundance of different classification levels. The taxonomy of each 16S rRNA gene sequence was analyzed by RDP Classifier (http://rdp.cme.msu.edu/) against the silva (SSU115)16S rRNA database using confidence threshold of 70%.

### 2.5 Statistical analysis

Comparisons of growth indicators were conducted by ANOVA. Differences were considered significant at p<0.05, and differences were considered to be extremely significant at p<0.01. Using Origin2021 software to draw graphics, all statistical analyses were carried out by SPSS 20.0 software. Structural equation modeling was established according to the previous report (Yang et al., 2023) and illustrated using R software.

## 3 Results and discussion

### 3.1 Tissue-specific responses of ginger plant for co-pollution resistance

#### 3.1.1 Physiological characteristics

Since the growth morphology of the crop intuitively reflects its response characteristics to external environmental factors, we first analysed the growth varations of ginger plants under different soil OFL and Cr treatment. From the morphology photo of ginger plant in rhizome expansion period (Figure 1A), it can be clearly observed that ginger plants under CrO100 treatment exhibited reduced branching, overall smaller body size, and more pronounced yellowing of leaf margins. After harvest, the height & fresh weight (FW) per ginger plant was measured as 60.17±2.3 cm & 317.70±6.9 g, 61.83±0.9 & 328.47±3.2 g, and 46.53±2.9 cm & 210.50±4.5 g under Cr, CrO1 and CrO100 treatment (Table 1), respectively. Regarding the growth of different organs, the root FW and rhizome FW under CrO100 treatment were found significantly reduced of 33.92% and 12.08% (P < 0.05), compared to Cr treatment. Besides, all plant growth indexes under CrO1 were slightly increased than Cr treatment, indicating a certain growth boosting effect of low-dose OFL introduction on Cr-stressed ginger.

**Figure 1.**
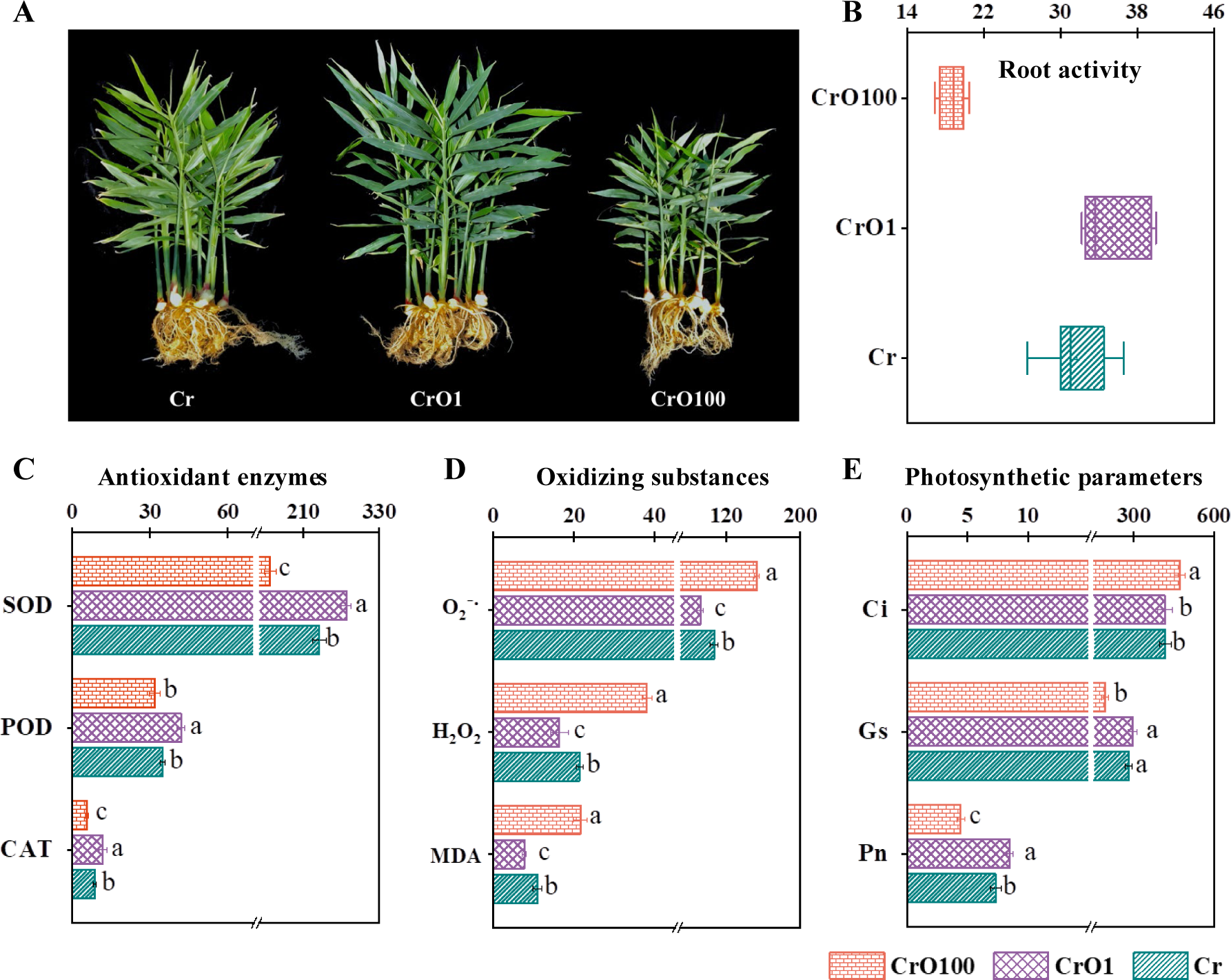
Physiological responses of ginger plants to OFL and Cr pollution. A. Morphology photo of ginger plant. B. Root activity (mg g^-1^(FW) h^-1^). C. Antioxidant enzyme activities including superoxide dismutase (SOD, units g^-1^), peroxidase (POD, ΔOD_470_ min^-1^ g^-1^), catalase (CAT, ΔOD_240_ min^-1^ g^-1^). D. Malondialdehyde (MDA, nmol g^-1^) and reactive oxygen species including hydrogen peroxide (H_2_O_2_, nmol g^-1^ min^-1^) and superoxide anion (O_2_^·-^, μmol g^-1^). E. Photosynthetic parameters including net photosynthetic rate (Pn, μmol m^-2^ s^-1^), stomatal conductance (Gs, mmol m^-2^ s^-1^) and intercellular CO_2_ concentration (Ci, μmol mol^-1^). Values followed with the same letter in the same figure was not significant at P = 0.05 (n=3), as well as in the later figures.

**Table 1.**
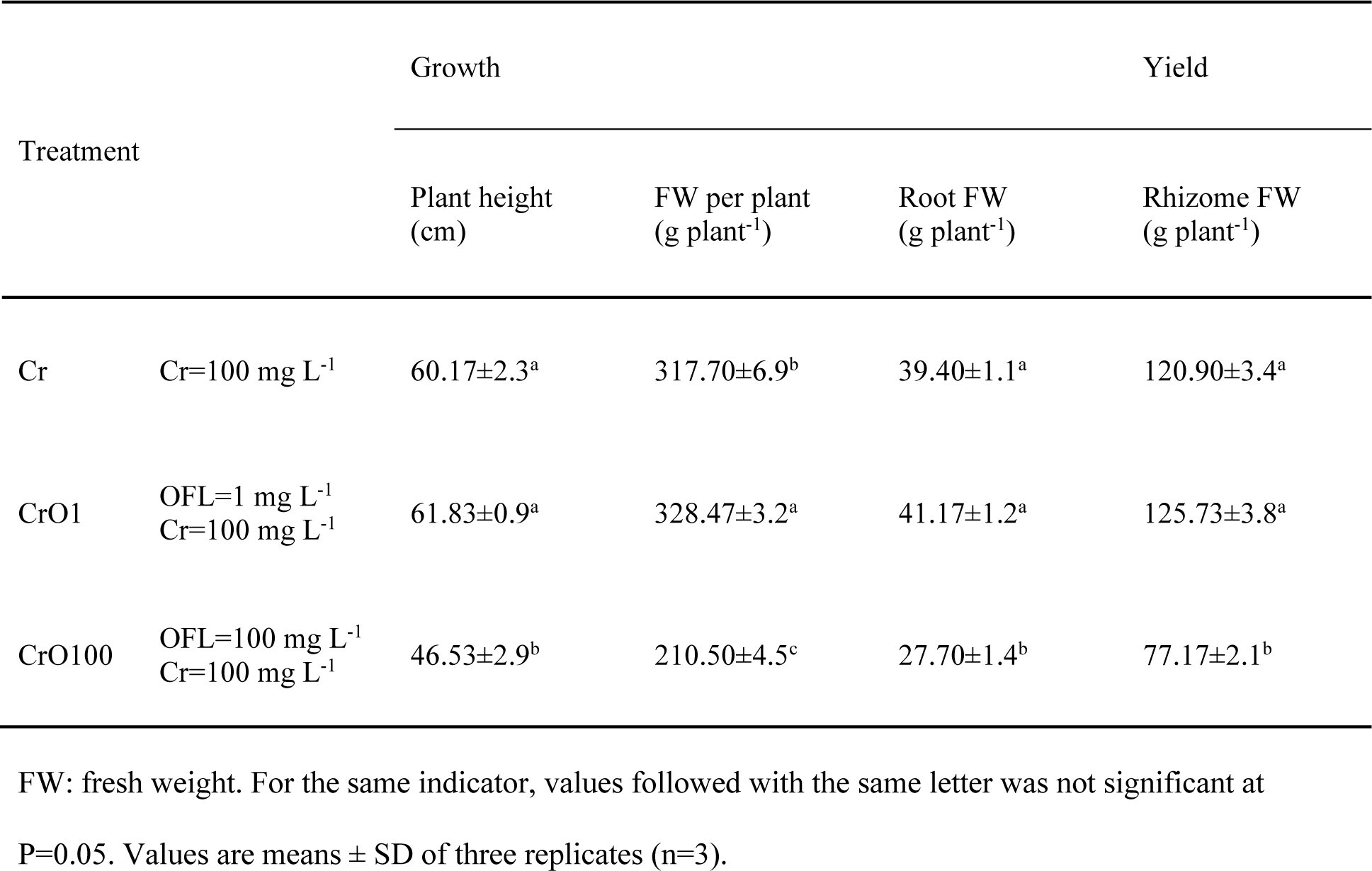
Interactive effects of soil ofloxacin and chromium on the growth and yield of ginger plants.

#### 3.1.2 Root system

In soil pollution conditions, roots sense OFL and/or Cr from soil at the first time. To be noted, low-dose OFL exhibted a hormesis on Cr-stressed ginger, i.e., exposure to low doses of a stressor can stimulate positive adaptive responses in organisms, resulting in enhanced resilience and improved physiological functions (Calabrese and Baldwin, 2003), which not only be capable of increasing the root growth but also enhancing the root activity (35.53 vs. 32.65 mg g^-1^(FW) h^-1^, Figure 1B). In contrast, for the roots of CrO100-treated plants, it was observed a significantly reduction in root activity than other treatments (P < 0.05), indicating a severe damage to the root system by the co-stress of Cr and high-dose OFL. The observed variation in root activity could be attributed to various factors. Earlier studies have reported the deleterious effects of Cr contamination in plant tissues, such as disrupt cellular processes and induce oxidative stress, which ultimately affects root development and vitality (Kushwaha and Singh, 2020). Here, our observations align with previous research that has explored the impact of Cr exposure, and also provides novel insights by investigating the combined effect of Cr and OFL, shedding light on potential antagonistic (CrO1) or synergistic (CrO100) interactions to ginger root systems between these stressors.

#### 3.1.3 Antioxidant system

Furthermore, such a considerable co-stress effect caused by CrO100 was reflected by the antioxidant enzyme activities in leaf system. As shown in Figure 1C, we can find that the activities of SOD, POD and CAT in the leaf system under CrO100 all decreased compared to single Cr treatment, in particular SOD and CAT reached a significant level of 43.36% and 53.19% reduction (P < 0.05). The involvement of antioxidative enzymes in mitigating oxidative stress is well-documented (Gill and Tuteja, 2010). Our findings suggested that the antioxidant enzyme defense mechanisms of leaves were compromised by the co-stress of Cr and high-dose OFL, potentially exacerbating the oxidative damage. Thus, the contents of MDA and reactive oxygen species (ROS) including H_2_O_2_ and O_2_^·-^ were further measured (Figure 1D). Here, the elevated levels of 97.98% MDA, 43.34% O_2_^·-^ and 78.63% H_2_O_2_ in ginger leaves, in response to CrO100 compared to Cr treatment further support the conclusion of ROS production and lipid peroxidation under overstress conditions (Gill and Tuteja, 2010). To be noted, the intriguing observations of increased antioxidant enzyme activities and decreased MDA and ROS levels under CrO1 treatment (Figure 1C&D) highlighted the possibility of a hormesis in the context of combined Cr and low-dose OFL stress in ginger leaf system. These findings indicated a potential adaptive response of Cr-stressed ginger, meaning that low-dose OFL can trigger an upregulation of antioxidative defense mechanisms to counteract oxidative damage. Meanwhile, the reduced MDA content suggests that low-dose OFL might trigger a protective response against lipid peroxidation, contributing to enhanced cellular integrity and function to Cr-stressed ginger (Morales and Munné-Bosch, 2019).

#### 3.1.4 Photosynthesis-fluorescence system

Photosynthesis in ginger plants is vital, converting light energy into chemical energy for growth. The measurements of photosynthetic parameters presented in Figure 1E showed CrO100-treated leaves had lower Pn, Gs, and higher Ci than other treatments (P < 0.05). This collective shift in these key parameters suggests severe leaf photosynthesis impairment due to Cr and high OFL co-stress. Here, Pn indicates the rate at which plants assimilate CO_2_ and produce organic compounds through photosynthesis. This decline in Pn (as 4.43 μmol m^-2^ s^-1^) is indicative of disrupted carbon fixation and energy production under CrO100, affecting plant growth and yield. Gs refers to the rate at which stomata, the tiny pores on leaves, open and allow water vapor to escape during transpiration. Diminished Gs (as 192.33 mmol m^-2^ s^-1^) indicates a reduction in water loss through transpiration. While this may initially appear as a positive response, decreased Gs can also limit the uptake of CO_2_ from the atmosphere, hindering photosynthesis (Lv et al., 2020). Ci reflects the CO_2_ concentration within the leaf’s internal spaces where photosynthesis occurs. Notably, the higher Ci (up to 471.33 μmol mol^-1^) with CrO100-stressed leaves suggests inefficient CO_2_ use for photosynthesis. Elevated Ci might signal uptake or fixation inefficiency due to disrupted stomatal regulation and biochemical processes (Todorenko et al., 2020). Such above findings align with the concept of stress synergy, where the combined effect of multiple stressors exceeds the sum of their individual effects, leading to heightened plant vulnerability and reduced fitness. Contrasting CrO100, CrO1 treatment shifted Pn, Gs, and Ci dynamics (Figure 1E), which highlights the potential of low-dose OFL to improve photosynthesis performances of Cr-stressed ginger, unveiling co-pollution interactions. Simply put, low-dose OFL maintained efficient transpiration and CO_2_ uptake and enhanced photosynthesis adaptively as reflected by the induced higher Pn and Gs via hormesis, echoing Lv et al.’s findings on low OFL-induced hormesis (Lv et al., 2020).

In addition, the visible color change in Fo, Fm and Fv/Fm (Figure 2A), with lighter hues indicating weaker photosynthetic function, further corroborated the photochemical inhibition effects under CrO100. According to Figure 2B, we can clearly observe a diminished PAR-ETR curve under CrO100, which further highlights compromised electron flow through the photosynthetic chain, potentially resulting from disruptions in photosystem II and associated electron transport processes (Xu et al., 2020a). Furthermore, Figure 2C reveals a substantial 19.69% and 11.21% reduction in φPSII and qP, respectivly, under CrO100 compaerd to Cr treatmen, along with a poor increase in NPQ. Here, reduced φPSII and qP indicate compromised photochemical efficiency and electron transport through photosystem I, possibly due to photosynthetic apparatus disruptions. Meanwhile, heightened NPQ suggests augmented energy dissipation, a safeguard against photodamage in stress conditions. Collectively, these findings signify significant leaf fluorescence system damage due to Cr and high-dose OFL co-stress. Conversely, under CrO1 treatment, a contrasting response in these fluorescence parameters were observed (Figure 2). The elevated φPSII and qP values, coupled with lower NPQ compared to other treatments, imply that low-dose OFL has a potentially amelioration in the photosystem’s fluorescence performance by improving photosynthetic apparatus integrity. In summary, the alterations in photosynthesis-fluorescence parameters underscore Cr and varying OFL doses’ intricate effects on the photosystem. The distinct responses observed under low-dose OFL provide evidence of potential amelioration, emphasizing the need to comprehend pollutant interactions to enhance plant resilience.The unique low-dose OFL responses provide evidence of potential amelioration, emphasizing the need to comprehend pollutant interactions to enhance plant resilience.

**Figure 2.**
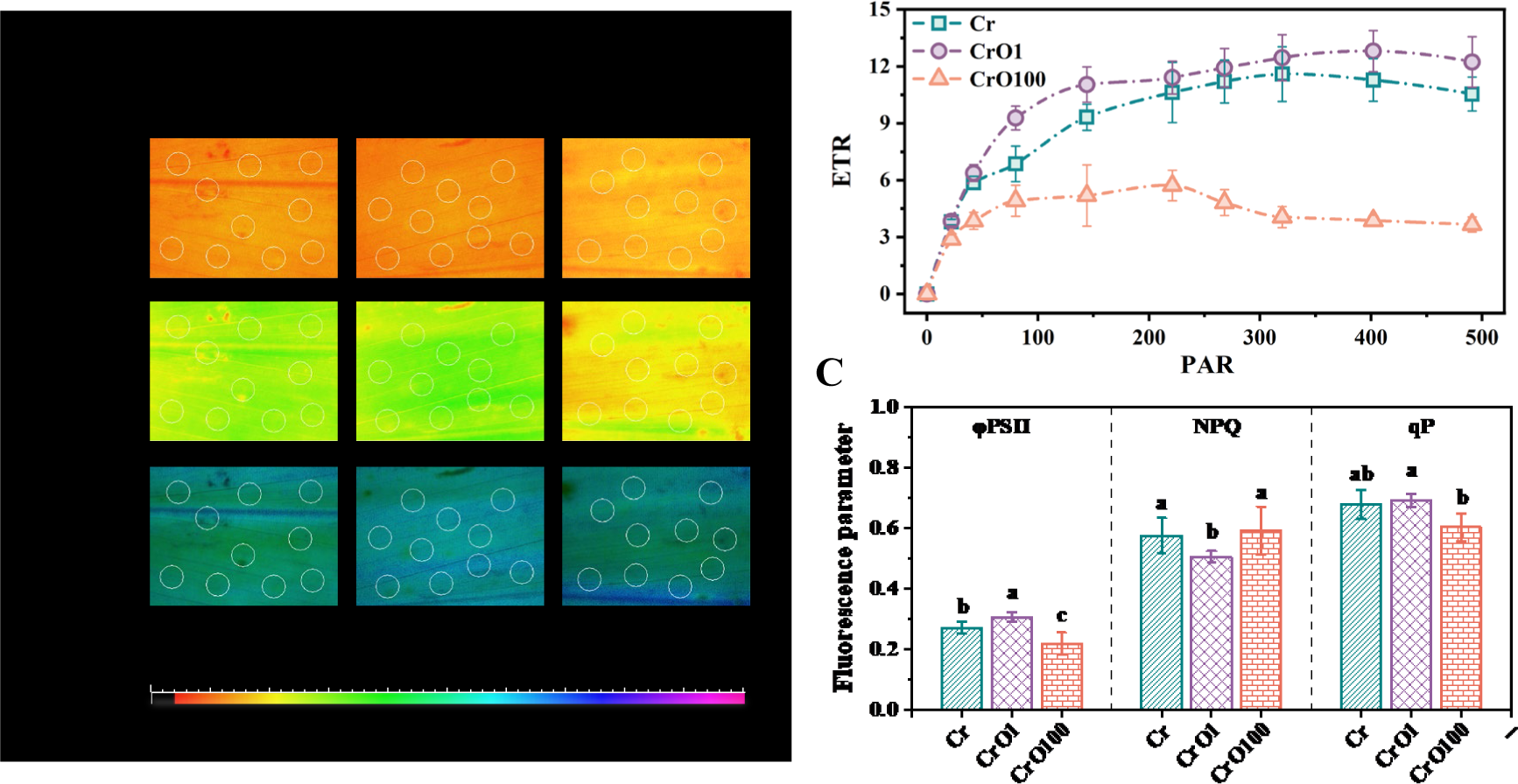
Chlorophyll fluorescence response of ginger leaves to OFL and Cr. A. Chlorophyll fluorescence images including initial fluorescence (Fo), maximal fluorescence (Fm) and the maximum photochemical efficiency (Fv/Fm). B. The photosynthetically active radiation-photosynthetic electron transfer rate (PAR-ETR) response curve of ginger leaves. C. The photochemical quantum efficiency of photosystem I (φPSII), non-photochemical quenching coefficient (NPQ) and the photochemical quenching coefficient (qP).

### 3.2 Responses of rhizosphere microbe for co-pollution resistance

#### 3.2.1 Rhizosphere microbial diversity and richness

High-throughput sequencing was conducted to determine the structure and composition of microbial populations among the bio-samples collected from the ginger rhizosphere. As listed in Table S2, the OTUs, diversity (*Shannon* and *Simpson*) and richness (*Chao* and *Ace*) index of microbial communities in different bio-samples all exhibited an order of CrO1 > CrO100 > Cr treatment. Moreover, the bacterial communities of these samples were clearly clustered into three groups, including Cr, CrO1 and CrO100 based on Principal Coordinates Analysis (PCoA) at the OTU level (Figure S1), while the sample points of CrO1 and CrO100 are closer together in the PCoA plot. Furthermore, the microbial ecological networks constructed based on OTUs revealed the intrinsic connections among different bio-samples (Figure 3A). Herein, a total of 3,665 OTUs (cluster 7) co-occurred in these three bio-samples. The unique OTUs in Cr, CrO1 and CrO100 were 129 (cluster 1), 234 (cluster 2) and 236 (cluster 3), respectively. These findings demonstrated a considerable shift in rhizosphere bacterial communities between the single pollution of Cr and the co-pollution of OFL and Cr, yet the differential clustering between CrO1 and CrO100 hinted that the outcome of such a shift varied across the OFL exposure dosage.

**Figure 3.**
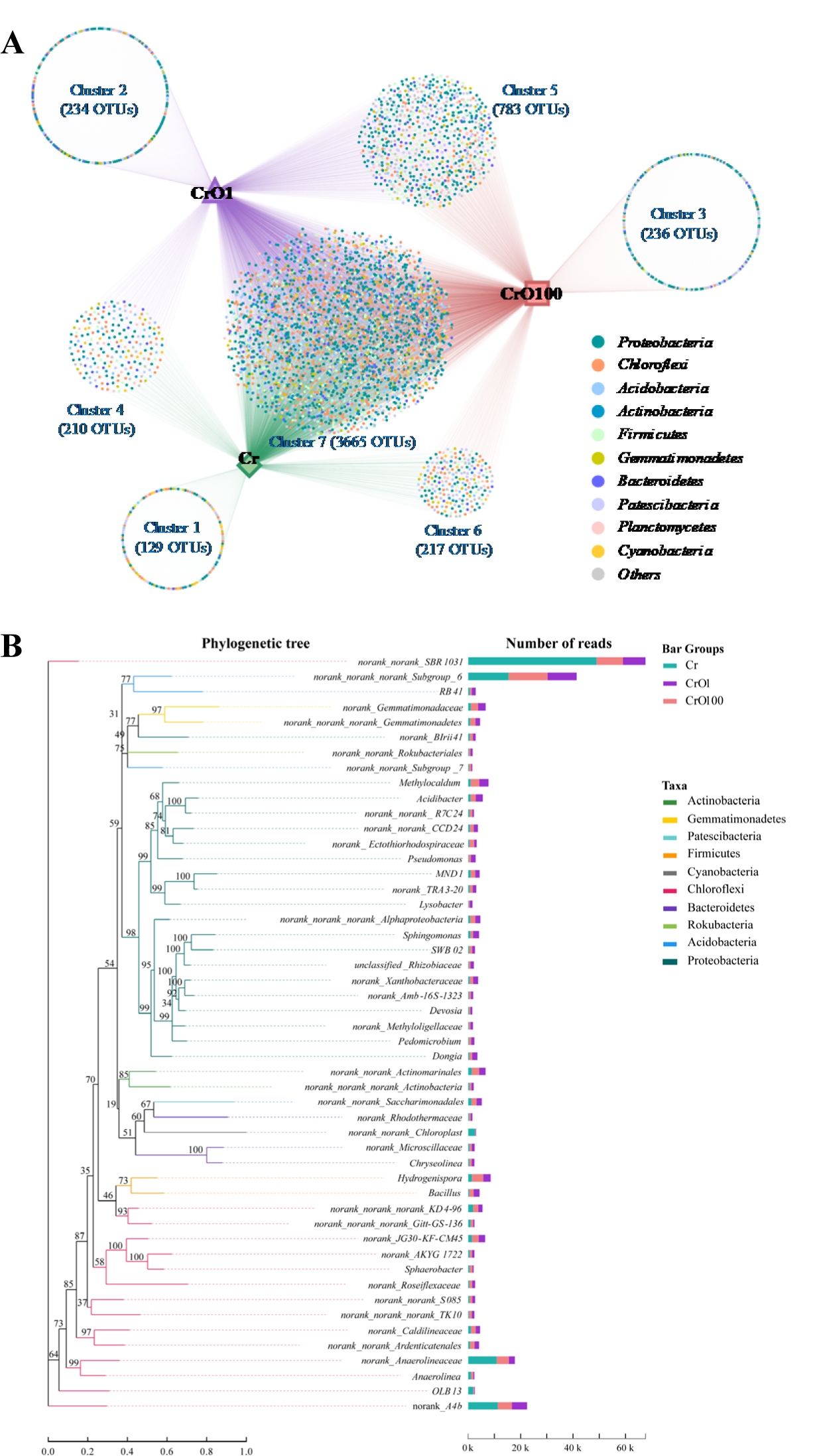
A. Network topology analysis for showing the presence and cross-correlation of microbial OTUs in different bio-samples. (Each circular node represents an OTU, where its different colors represent different phyla (Top 10) classifications; thicker connecting lines indicate higher relative abundance (weight) of the corresponding OTU; the numbers s in parentheses represent the amounts of intersections or unique OTUs in each sample region. B. Phylogenetic tree illustrating systematic evolutionary relationships on genus level. (Branches represent species categories and are color-coded by higher taxonomic levels. Branch length indicates evolutionary distance, reflecting species differences. Bar chart displays species’ Read proportions across various groups).

#### 3.2.2 Rhizosphere microbial composition and evolution

To gain insight into the association between the rhizosphere bacterial community structures and pollution exposures, the beacterial phlya and genera were respectively identified according to taxonomic affiliation. As shown in Figure S2A, the original dominant phyla under different treatments are similar and not varied during community evolution, including Chloroflexi (57.38% vs. 22.40% vs. 39.97%, indicate Cr vs. CrO1 vs. CrO100, hereinafter the same), Proteobacteria (13.68% vs. 37.40% vs. 30.82%) and Acidobacteria (12.67% vs. 10.74% vs. 13.09%). To be noted, the bacterial aboundance variations among different treatments indicated that the presence of OFL alongside Cr might create a different selective pressure that favors Proteobacteria over Chloroflexi. In fact, Proteobacteria is known for its exceptional metabolic versatility and adaptability to thrive under varying environmental conditions, as well as its high intrinsic resistance to antibiotics (Chevalier et al., 2017). Furthermore, the phylogenetic tree depicted in Figure 3B illustrates systematic evolutionary relationships at the genus level (Top 50). Here, it is evident that the representative genera *norank_f__norank_o__SBR1031* (33.21% vs. 5.87% vs. 6.78%, Figure S2B) and *norank_f__A4b* (7.66% vs. 3.95% vs. 3.63%) within the Chloroflexi exhibit a significant decrease in abundance under co-stress conditions. Similarly, previous studies have documented a noticeable reduction of these two genera under antibiotic selection pressure (Sardar et al., 2021; Song et al., 2020). Additionally, it is noteworthy that a total of 20 genera belonging to the Proteobacteria exhibit a distinct trend of proliferation under co-exposure to OFL and Cr, such as *Methylocaldum* (0.62% vs. 2.40% vs. 2026%), *Acidibacter* (0.60% vs. 1.86% vs. 1.35%), *norank_f__Ectothiorhodospiraceae* (0.32% vs. 0.64% vs. 1.24%), and *Pseudomonas* (0.13% vs. 1.25% vs. 0.58%). This could be attributed to the selective pressure exerted by OFL on rhizosphere bacterial communities. As reported, *Methylocaldum*, *Acidibacter* and *Pseudomonas* might possess antibiotic resistance or degradation functions (Chevalier et al., 2017; Man et al., 2020; Wan et al., 2021).

### 3.3 Interactive characteristics of microbial-crop for co-pollution resistance

#### 3.3.1 Root exudates variations

Ginger root exudates were investigated to understanding interactive characteristics of microbial-crop system. According to PARAFAC analysis results, five fluorescent components of root exudates were identified and summerized in Table S3, which could be attributed to visible fulvic-like (C1 and C2) and humic-like (C3, C4 and C5). This characterization is consistent with previous research that identified similar fluorescent components in plant root exudates exposed to heavy metal or antibiotic stress (Chen et al., 2018; Xu et al., 2023). Further, the maximum fluorescence intensity (Fmax) was employed to assess variations in root exudates under different pollution scenarios. Higher Fmax values were indicative of greater content of specific exudate components. As shown in Figure S3, the Fmax exhibited a slight increase (minly humic-like exudates) under CrO1 compared to single Cr treatment, indicating a potential mitigating or even enhancing effect of low-dose OFL on root exudate patterns. As known, the functional groups of humic-like compounds, such as carboxylic and phenolic moieties, can interact with diverse compounds. Studies have indicated that humic-like can sorb antibiotics like OFL as well as heavy metals like Cr, influencing their transport and bioavailability in soil systems (Aldmour et al., 2019; Guillossou et al., 2021). However, the Fmax under CrO100 was significantly reduced than other treatments, which could be attributed to the pronounced adverse impacts of the co-stress of Cr and high-dose OFL on the structure and function of root system as montioned in Section 3.1.

#### 3.3.2 Antibiotic resistance genes enrichment and co-occurrence with microbes

Microbial community functional predictions were performed using PICRUSt (version 1.0.0-dev) against the Kyoto Encyclopedia of Genes and Genomes (KEGG), followed by an in-depth analysis of pathogens associated with human diseases. As depicted in Figure 4B, six disease-relevant pathogens across different treatments, accounting for approximately 6∼7‰ of the total rhizosphere bacteria, followed the order of CrO1 (6.89‰) > CrO100 (6.67‰) > Cr (6.60‰). These proportions closely resemble the reported distribution of pathogenic bacteria in natural water bodies (<10‰), such as Ebinur Lake Basin (Wang et al., 2021) and Tiaoxi River (Zheng et al., 2017). Notably, the disease resistance: antimicrobial-related pathogens increased by 65.54% and 66.76% in CrO1 and CrO100 treatments respectively compared to Cr treatment, suggesting that the presence of OFL induced an elevation in drug resistance within the microbial community, thereby posing a higher risk of ARGs dissemination. Similarly, prior studies have affirmed the connection between pathogens and antibiotic resistance genes (ARGs) in greenhouse soils, river waters, and lake systems (Fan et al., 2018; Wang et al., 2021; Zheng et al., 2017), signifying the potential threat they pose to human health.

**Figure 4.**
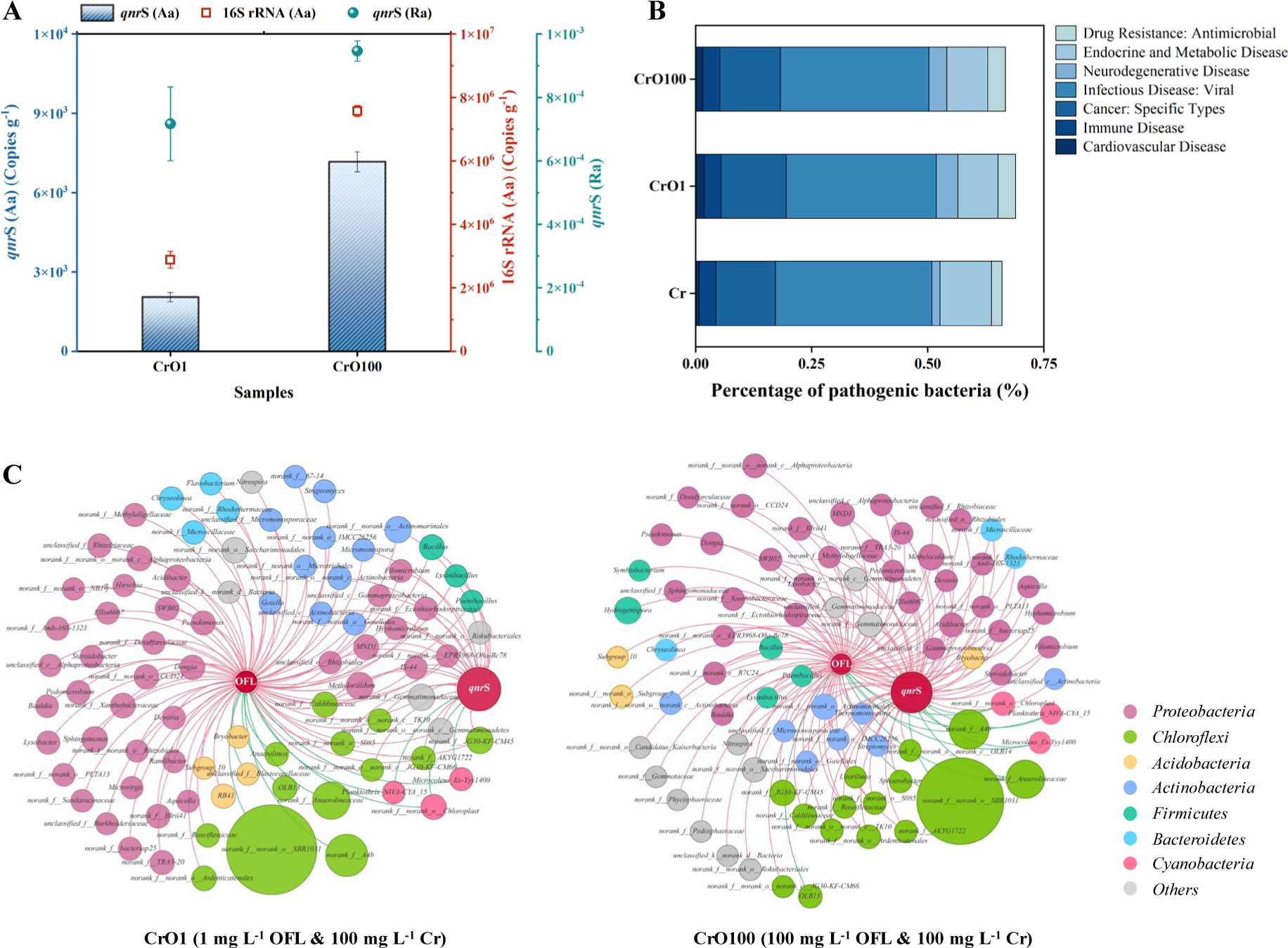
A. Absolute abundance (Aa) and relative abundance (Ra) of typical antibiotic resistance gene (*qnr*S) detected in ginger rhizomes. B. Percentage of pathogenic bacteria in different rhizosphere microbial samples. C. Co-occurrences of OFL, *qnr*S and their potential host bacteria under co-pollution. Nodes stand for bacterial species at genus, OFL and ARG subtypes. Different node colors represent different phyla. A connection (i. e., edge) represents a significant pairwise correlation (P < 0.05), and a red edge represents a positive correlation while a green edge represents a negative correlation.

Moreover, the typical ARG (*qnr*S) associated with quinolone antibiotics in FOL- and Cr- co-polluted rhizomes (edible organ) was investigated to understand its potential transmission risk within the food chain. The relative abundance of *qnr*S detected in CrO100 was 1.32 times higher than that in CrO1 (7.17×10^-4^ copies/16S rRNA vs. 9.46×10^-4^ copies/16S rRNA), indicating the stimulation of high-dose OFL on qnrS expression in ginger rhizomes. Furthermore, given the demonstrated ability of environmental heavy metals to facilitate the horizontal transfer of ARGs (de Voogt, 2017), the presence of Cr might also promote the enrichment of *qnr*S within plant organs to some extent. Furthermore, network analysis was employed to unveil co-occurrence patterns between environmental factors and rhizosphere bacteria (Figure 4C). Notably, under CrO1 co-pollution, 75 and 26 genera were positively correlated with OFL and *qnr*S respectively, while under CrO100 co-pollution, 73 and 55 genera respectively correlated positively with OFL and *qnr*S. This finding implies that the selection stress induced by high-dose OFL significantly increased the potential host bacteria harboring the *qnr*S gene. The dominant ARG-associated genera mainly came from Proteobacteria, a taxonomic group known for high ARG abundance in microbial communities, particularly in soils (Bahram et al., 2018). More importantly, the proliferation of potential hosts in the context of co-pollution may link to the disease resistance: antimicrobial-related pathogens (Figure 4B) and dirve higher ARG abundances into ginger rhizomes (Figure 4A), as alterations in host composition play a primary role in ARG alterations (Li et al., 2019).

#### 3.3.3 Possible interactive mechanism

By collecting the aforementioned data, we employed structural equation modeling to establish quantitative correlations between core indices and the crop-microbe system. In Figure 5A, a negative correlation between latent variables of soil & crop was observed, with a standardized path coefficient (SPC) of −0.82 (p < 0.05, hereinafter the same). Additionally, crop and its measurement factor (MF) of reactive oxygen species (ROS) displayed a negative correlation (SPC ∼1). These results are in accordance with previous findings suggesting that soil pollution may diminish crop health and productivity, primarily through elevated ROS levels (Lv et al., 2020; Zhang et al., 2023). Furthermore, clear positive correlations were evident between factors of soil & microbe, as well as microbe & crop, with SPCs of 0.36 and 0.33 (p < 0.005), respectively. Moreover, crop and its MFs of diversity (Shannon), richness (Chao), and pathogenic bacteria (Antimicrobial) exhibited positive correlations. This aligns with our aforementioned analysis, indicating that under co-pollution (i.e., CrO1, CrO100), OFL induced greater diversity, richness, and disease resistance: antimicrobial-related pathogens compared to single Cr pollution.

**Figure 5.**
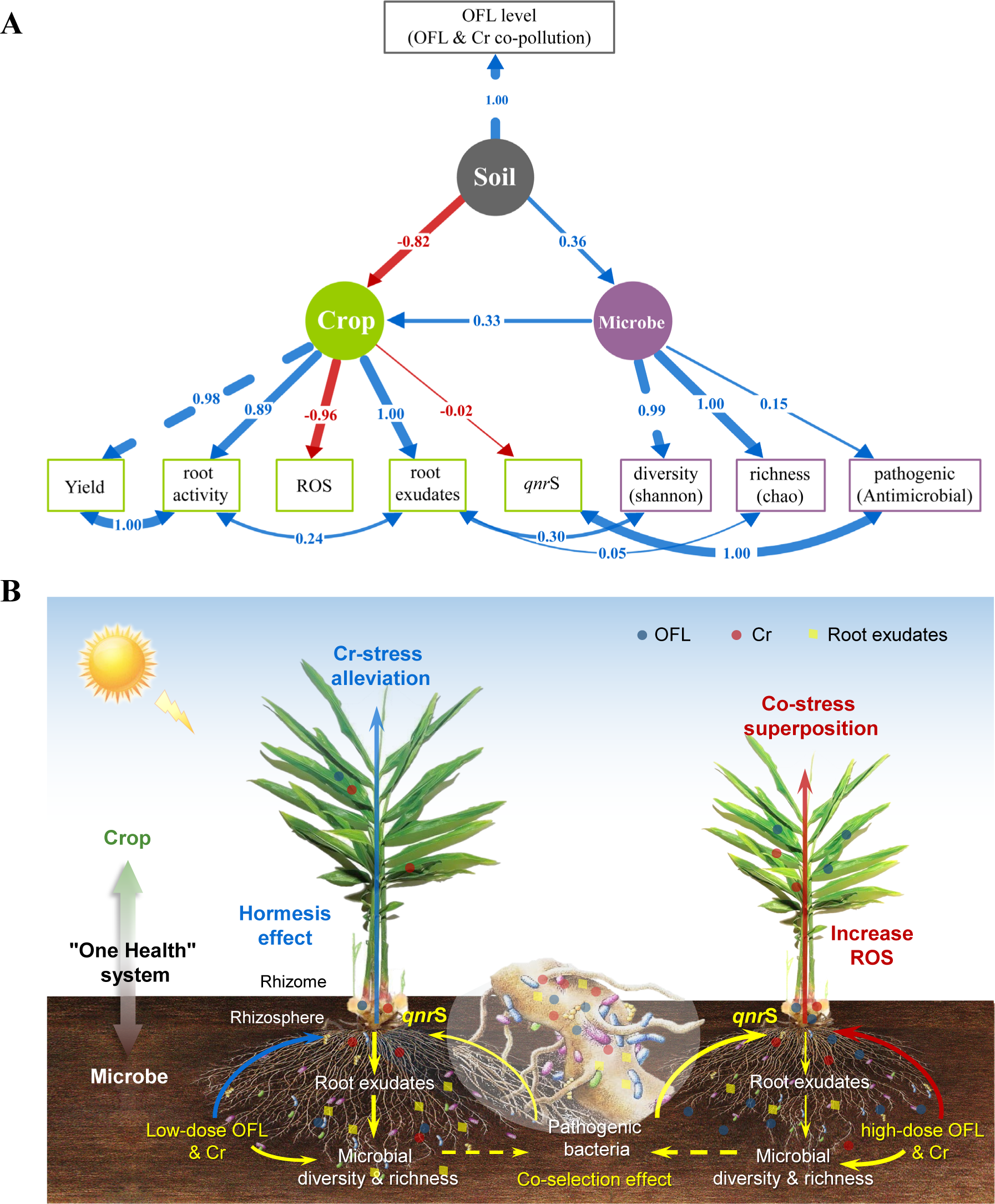
(A) Structural equation modeling for the quantitative correlations between core index of crop-microbe system to soil OFL and Cr co-pollution. Standardized coefficients were used as the path effects. Solid lines represented the significant path (p < 0.05) and dash lines represented non-significant paths. (B) Schematic diagram of the tissue-specific responses and interactive characteristics of crop-microbe “One Health” system to OFL and Cr co-pollution.

Root exudates are known to play essential roles in supporting plant growth and interacting with physicochemical and biological factors in the rhizosphere (Chai and Schachtman, 2022). In this study, root exudates positively correlated with root activity (SPC ∼0.24), microbial diversity (SPC ∼0.30), and richness (SPC ∼0.05), providing solid evidence for a positive interaction between the crop and microbe. In fact, the pronounced positive correlation (SPC ∼1) between *qnr*S and pathogenic bacteria (*Antimicrobi*al) warrants particular attention due to the potential negative impact of *qnr*S enrichment in edible crop organs on crop health (SPC ∼-0.02).

In summary, a schematic diagram illustrating the tissue-specific responses and interactive characteristics of the crop-microbe “One Health” system to OFL and Cr co-pollution is presented in Figure 5B. In brief, under soil co-pollution conditions, low-dose OFL exhibits a hormesis effect on crop growth, enhancing physiological and metabolic functions (e.g., root activity, antioxidant enzyme activities, photosynthesis-fluorescence) of crop tissue systems, effectively counteracting Cr-induced ROS stress. However, it’s important to note that this hormesis effect has limitations. However, such a hormesis effect has certain limitations. When exposed to high-dose OFL, the superposition effects of co-pollution lead to severe damage to crop growth and metabolic functions, inducing excessive ROS and ultimately resulting in reduced crop yield. Additionally, rhizosphere microbial diversity and richness are vital potential factors in driving crop tolerance to soil pollution stress, which might be attributed to the increased capacity of rhizosphere microbes to facilitate the biotransformation of pollutants (Aldmour et al., 2019; Guillossou et al., 2021). To be noted, the presence of OFL, especially high-dose OFL, also triggers the proliferation of antimicrobial-related pathogens. This phenomenon may facilitate the transfer and buildup of ARGs within edible crop organs, subsequently propagating along the food chain and exacerbating associated risks.

## 4 Conclusion

This study provides significant insights into the complex interplay between OFL and Cr co-pollution within the crop-microbe system. By examining the tissue-specific responses and interactive characteristics, we have addressed a critical gap in previous research. The results indicate that low-dose OFL induces hormesis effects on ginger plant growth and physiological functions, enhancing root activity, antioxidant defenses, and photosynthesis efficiency in response to Cr-induced stress. However, high-dose OFL exacerbates oxidative damage and inhibits plant growth. Rhizosphere microbial diversity and composition are influenced by co-pollution, with Proteobacteria proliferating under OFL influence. Additionally, the enrichment of *qnr*S gene and potential pathogenic bacteria raises concerns about environmental and food chain risks. More importantly, this study quantitatively establishes correlations within the soil-crop-microbe system through a structural equation model, highlighting the interconnected nature of these interactions. This research contributes to advancing our understanding of the soil co-pollution effects within crop-microbe “One-Health” system, thus laying the foundation for informed decision-making in sustainable agricultural and environmental management practices.

## Acknowledgement

This research was supported by the Agricultural Fine Variety Project in Shandong Province of China (Grant No. 2020LZGC006), China Agriculture Research System (Grant No. CARS-24-A-09), National Natural Science Foundation of China (Grant No. 31972399), Natural Science Foundation of Shandong Province (Grant No. ZR2022QC009) and National Natural Science Foundation of China (Grant No. 52222003).

